# Secondary single-cell transcriptomic analysis reveals common molecular signatures of cerebrovascular injury between traumatic brain injury and aging

**DOI:** 10.1101/2020.06.29.178855

**Authors:** Xinying Guo, Bangyan Zhang, Fernando Gomez-Pinilla, Fan Gao, Zhen Zhao

## Abstract

Cerebrovascular injury is a common pathological feature of a spectrum of neurological disorders including traumatic brain injury (TBI), stroke, Alzheimer’s disease (AD), as well as aging. Vascular manifestations among these conditions are similar indeed, including the breakdown of the blood-brain barrier (BBB). However, whether there is a common molecular mechanism underlying the vascular changes among these conditions remains elusive. Here, we report secondary transcriptomic analysis on cerebrovascular cells based single-cell RNA-seq datasets of mouse models of mild TBI and aging, with a focus on endothelial cells and pericytes. We identify several molecular signatures commonly found between mTBI and aging vasculature, including *Adamts1, Rpl23a, Tmem252, Car4, Serpine2*, and *Ndnf* in endothelial cells, and *Rps29* and *Sepp1* in pericytes. These markers may represent the shared endophenotype of microvascular injury and be considered as cerebrovascular injury responsive genes. Additionally, pathway analysis on differentially expressed genes demonstrated alterations in common pathways between mTBI and aging, including vascular development and extracellular matrix pathways in endothelial cells. Hence, our analysis suggests that cerebrovascular injury triggered by different neurological conditions may share common molecular signatures, which may only be detected at the single-cell transcriptome level.

## Introduction

Traumatic brain injury (TBI) is a leading cause of death and disability, particularly in young adults thereby resulting in much greater loss of productivity compared to other conditions(Sandsmark et al., 2019). It is also considered as the most robust environmental risk factor for Alzheimer’s disease (AD), and it is highly prevalent during military service and contact sports and doubles the risk of developing AD and dementia (Sibener et al., 2014). Mechanistically, TBI triggers multiple neurodegenerative cascades, including axonal damage and excitatory toxicity in neurons, inflammatory responses in microglia and astrocyte, and cerebral microvascular impairment such as local edema, blood-flow reduction, and breakdown of the blood-brain barrier (BBB) (Jullienne et al., 2016; Ramos-Cejudo et al., 2018; Sandsmark et al., 2019). However, the underlying molecular mechanism of microvessel injury in TBI, particularly the mild cases, remains elusive.

The cerebrovasculature is constituted by endothelial cells (EC), mural cells including pericytes (PC) and vascular smooth muscle cells (VSMC), and vessel-associated cell types including astrocytes and fibroblast cells (Vanlandewijck et al., 2018). The BBB is a central and built-in function of cerebral vessels, which serves as a unique interface between the circulation and central nervous system (CNS). Histological and neuroimaging assessments have demonstrated that microvessel injury with BBB dysfunction is common in TBI. It can occur during the acute phase of TBI and last for years in patients(Wilson et al., 2017), resulting in long-term adverse consequences of brain functions in nearly 50% of the patients (Hay et al., 2015). This microvascular endophenotype was well recapitulated in periclinal animal models, which demonstrated that TBI induces endothelial dysfunction (Villalba et al., 2017) and disrupts endothelial-pericyte crosstalk (Bhowmick et al., 2019), resulting in cerebral blood flow reduction and tissue hypoxia(Foley et al., 2013), leukocyte infiltration, gliosis and activation of neuroinflammation(Simon et al., 2017).

Cerebrovascular injury and BBB impairment are also commonly found in many neurological conditions, including but not limited to TBI, stroke, AD, amyotrophic lateral sclerosis (ALS), multiple sclerosis (MS) as well as in aging (Sweeney et al., 2019). Cerebrovascular injury can lead to the disruption of BBB integrity, including influx of blood-derived neurotoxin and collapse of microcirculation, and alteration of BBB function, including EC and mural cells dysfunction, and eventually accelerate neuronal damage, injury and degeneration (Sweeney et al., 2019; Zhao et al., 2015). It has been reported that cerebrovascular injuries facilitate the production of toxic substances and perivascular accumulation. These neurotoxins further cause BBB disruption, driving a destructive feedback loop(Sweeney et al., 2018; Zlokovic, 2011). Therefore, BBB dysfunction is now increasingly appreciated as an early event and an important driving force of neuropathological changes that participate in neurodegenerative diseases. Although cerebrovascular injury is important in many diseases, many questions are still elusive. Here, we established a workflow to accurately transfer single-cell RNA sequencing (scRNA-seq) information across different datasets, and determined the common molecular signatures underlying the microvascular injury at least in mTBI and aging. We further discuss the potential mechanisms of how common molecular signatures induce pathophysiological changes in cerebrovascular injury and increase the risks of developing multiple neurodegenerative diseases after mTBI.

## Materials and Method

### ScRNA-seq data for mTBI and aging sample

For the mTBI scRNA-seq dataset (Arneson et al., 2018), we obtained the cell count matrix from Gene Expression Omnibus (GEO) with the series record GSE101901. The data represents the expression level of 20728 genes in 3853 and 4544 hippocampal cells from mice suffered mild fluid percussion injury (FPI) or Sham surgery (n=3 per group), respectively. The hippocampal tissue was harvested 24 h after procedures for Drop-seq. The scRNA-seq libraries were prepared with Nextera DNA Library Preparation kit (Illumina, San Diego, CA, USA) and sequencing was performed on an Illumina HiSeq 2500 (Illumina, San Diego, CA, USA) instruments.

For the aging scRNA-seq dataset (Ximerakis et al., 2019), we obtained the cell count matrix from GEO with the series record GSE129788. The data represents the expression level of 14699 genes in 16028 and 21041 brain cells from young mice or old mice (n=8 per group), respectively. Young male C57BL/6J(B6) mice at the age of 2-3 months and old male mice at the age of 21-22 months were used. The whole brain tissue was harvested at the same time for 10X Genomics. The scRNA-seq libraries were prepared with Chromium Single Cell 3’ Library & Gel Bead kit v2 and i7 Multiplex kit (10X Genomics) and sequencing was performed on a NextSeq 500 instrument (Illumina).

### Data preprocessing

The data processing of the scRNA-seq data was performed with the Seurat Package (v.3.1.5) in R (v.3.6.1). The basic scRNA-seq analysis was run using the pipeline provided by Seurat Tutorial (https://satijalab.org/seurat/v3.0/immune_alignment.html) as of June 24, 2019. In general, we set up the Seurat objects from different groups in experiments for normalizing the count data present in the assay. This achieves log-normalization of all datasets with a size factor of 10,000 transcripts per cell. For different Seurat objects, FindVariableFeatures() function was used to identify outlier genes on a ‘mean variability plot’ for each object. The nFeatures parameter is 2000 as the default for the selection method called ‘vst’. These resulted genes serve to illustrate priority for further analysis.

### Integration of scRNA-seq data

A new integration method was used to integrate the Seurat Object from different groups(Stuart et al., 2019). Using the FindIntegrationAnchors() function, we found a set of pairwise correspondences between individual cells. We used the default dimensions (1:20) to be used for the canonical correlation analysis, which assumes that cells originate from a common biological state will be matched based on the first 20 dimensions. These anchors are used for downstream integration of the objects. We used the IntegrateData() function with the previously computed anchor set as a parameter to integrate the two Seurat object. The default dims parameter (1:20) was used for the anchor-weighting procedure.

### Data processing

The integrated dataset on all cells was used to scale and center the genes. First, principal component analysis (PCA) was used for linear dimensionality reduction with default settings to compute and store 30 principal components. Then, the top 20 principal components were used as the input for the Uniform Manifold Approximation and Projection (UMAP) dimensional reduction. We identified clusters of cells by a shared nearest neighbor (SNN) modularity optimization-based clustering algorithm. The algorithm first calculated k-nearest neighbors and computed the k-NN graph, and then optimizes the modularity function to determine clusters.

### Determination of cell-type identity

To determine the cell type, we identified marker genes for each cluster, by performing differential gene expression analysis for each cluster against all other clusters independently, using a Wilcoxon Rank Sum test. The results of each differential analysis were ordered based on log fold change of the average expression (genes with the highest fold-change receiving the lowest rank number). Top_n markers indicated the top n ranking maker genes. To assess statistical significance, we generated adjusted P-value by using Bonferroni correction. Cell-type specific marker genes to determine cell identity were based on previous reports (Saunders et al., 2018), include, but are not limited to *Snap25* for all neurons, *Gpc5* for astrocytes, *Mog* for oligodendrocytes, *Vcan* for oligodendrocyte precursor cells (OligoPC), *Cldn5* and *Rgs5* for vascular cells, *Ctss* for microglia. To identify the subtype of neurons and vascular cell types, we compared their signature to known hippocampal cell types derived from large scale scRNA-seq in mice(La Manno et al., 2016),(Zeisel et al., 2015), and identified *Spink8* for neurons in Cornu Amonis (CA), *C1ql2* for neurons in dentate gyrus (DG), *Gad2* for interneurons, *Nhlh2* for Caja-Retzius neurons, *Kcnj13* for choroid plexus cells, *Dcn* for fibroblast-like cells and *Tmem212* for ependymal cells.

### Secondary clustering for vascular cells

The cell cluster which contains canonical EC markers (*Cldn5, Pecam1, Kdr, Flt1*, and *Icam2*), PC markers (*Kcnj8, Pdgfrb, Rgs5, Vtn*, and *Abcc9*), and VSMC markers (*Acta2, Tagln, Myh11, Myl9*, and *Mylk*) is considered as vascular cell cluster. For the vascular cell cluster, we re-clustered the subcategorized cell types following the same strategy which were shown above. In general, we took vascular cells for scaling, performing PCA and UMAP, clustering, and identification of cell types. In the mTBI dataset, we identified 468 vascular cells (236 from sham group and 232 from mTBI group), while in the aging dataset, we identified 3459 vascular cells (1450 from young mice and 2009 from aging mice). In the secondary analysis for vascular cells, most of EC, PC, VSMC, and astrocyte makers are consistent with previous reports (Vanlandewijck et al., 2018; Ximerakis et al., 2019).

### Identification of DEGs and pathway analysis

After the determination of cellular identities, we performed differential expression genes based on the non-parametric Wilcoxon rank-sum test and generated statistical perimeters including P_value, avg_logFC, pct.1, and pct.2 (pct is the percentage of cells where the genes are detected in each group). The genes with at least a 0.25 log fold change between different conditions and P-value < 0.01 were considered as Differential Expression Genes (DEGs). The DEGs were used in downstream pathway enrichment analyses. Enrichment of pathways from KEGG, GO Biological Process, GO Molecular Function, and GO Cellular Components was assessed with the hypergeometric test and the Benjamini-Hochberg p-value correction algorithm to identify ontology terms.

### Meta-analysis by Metascape

Using the Metascape online tool (http://metascape.org), we performed functional enrichment analysis of multiple gene lists from different datasets. Enrichment of pathways from KEGG, GO Biological Process, GO Molecular Function, and GO Cellular Components was analyzed by Metascape. The terms with P-value < 0.01, minimum counts of 3, and enrichment factors of > 1.5 would be considered.

## Result

### Secondary analysis of cerebrovascular changes in mTBI scRNA-seq dataset

To understand the molecular signature of mTBI-induced microvascular injury, we analyzed the scRNA-seq data of hippocampal cells from mTBI and sham animals(Arneson et al., 2018). The processed cell count matrix from the study of scRNA-seq of mouse hippocampus tissues in mTBI was collected for the secondary analysis using the R Seurat Package. Cell count matrix from sham and mTBI groups were integrated to identify ‘anchors’ across diverse single-cell data types (Stuart et al., 2019), and the top 20 principal components from PCA analysis were used to generate UMAP (**Fig. 1A**). In the UMAP space, 12 cell types were separated into clusters (**Fig. 1B**) and identified using specific genetic markers (**Sup. Table1**) and their distinct cluster-specific expression patterns (**Sup. Fig. 1A**), such as *Mog* for oligodendrocytes, *Ctss* for microglia, *Vcan* for oligodendrocyte precursor cells (OligoPC), *Gpc5* for astrocytes. To validate the reliability of our analysis, we chose the top 3 gene markers in each cell type and compared their expression patterns with DropViz(La Manno et al., 2016), and found that the cell type specific genes in our dataset are largely consistent with their results from 690,000 cells in adult mouse brain (**Sup. Fig. 1B)**. Next, we separated cells between the mTBI mice and their controls in each cell type (**Fig. 1C**). The abundance of most cell types was largely consistent between sham and mTBI groups (**Fig. 1D**), except for injury-associated decline of all three types of neurons in CA and DG after mTBI.

**Figure 1.**
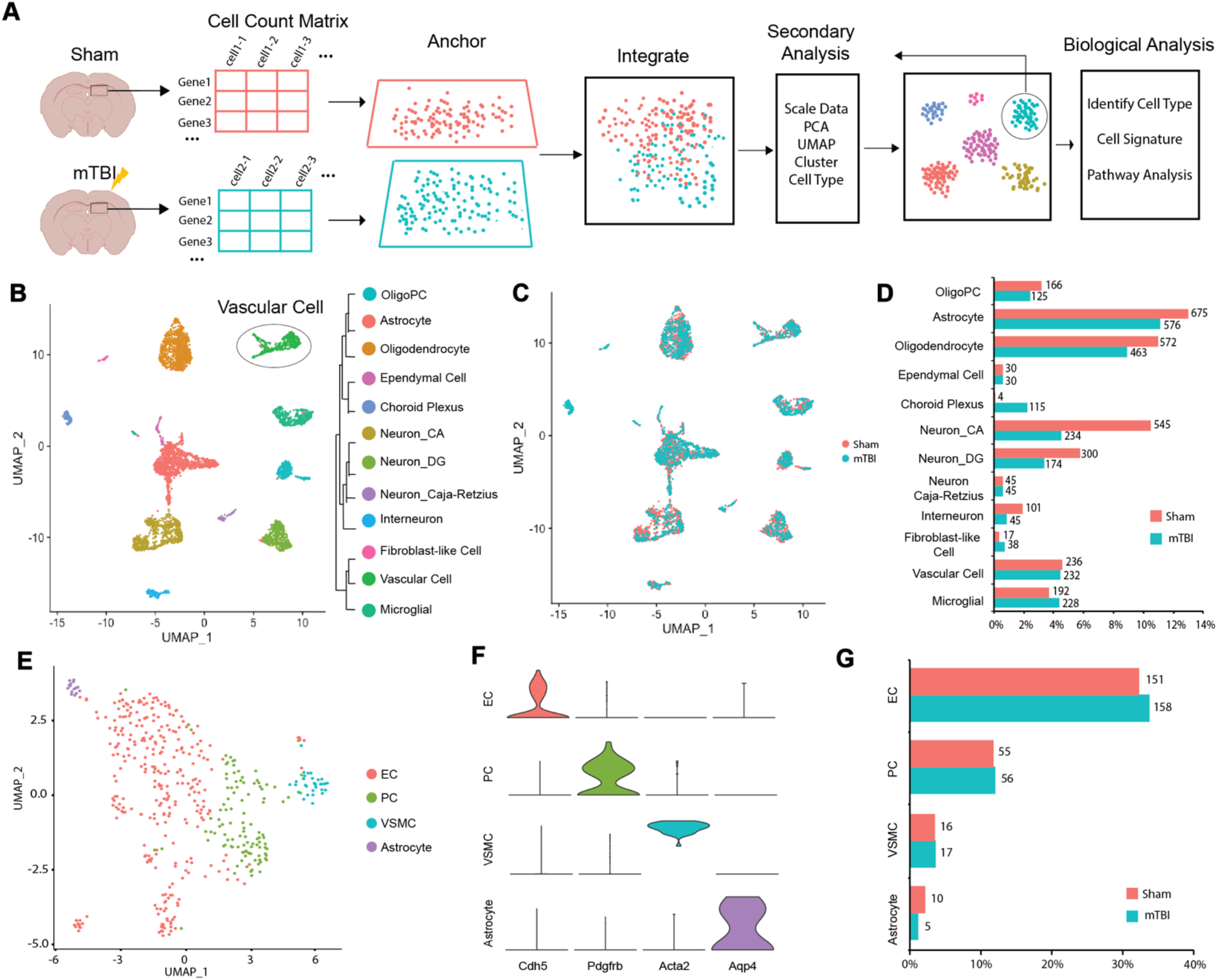
Secondary analysis of cerebrovascular changes in mTBI scRNA-seq dataset. **A**, Schematic of data collection and analysis workflow. **B**, UMAP of 5,192 single cell transcriptomes (2,883 from of sham group and 2,309 from mTBI group). Each dot was color coded and annotated based on their transcriptional profile identities. Vascular cells are marked by black line. OligoPC: Oligodendrocyte precursor cell; CA: Cornu amonis; DG: Dentate gyrus. **C**, UMAP showing an individual cell, with sham group in peach and mTBI group in turquoise. **D**, Bar plot showing the percentage of cells associated with each cell type in both sham and mTBI group. The number besides the bar shows cell number in each cell type of sham and mTBI group. Peach bar represents sham group and turquoise bar represents mTBI group. **E**, UMAP visualization of 468 vascular cell transcriptomes (236 from of sham group and 232 from mTBI group). UMAP showing cell clusters which were color coded and annotated based on their transcriptional profile identities. EC: Endothelial cell; PC: Pericyte; VSMC: Vascular Smooth Muscle Cell. **F**, Violin plots showing gene markers that distinguish across vasculature cells. Genes are colored by cell types. **G**,Bar plot showing the percentage of cells associated with EC, PC, VSMC, and Astrocyte in both sham and mTBI group. The number besides the bar shows cell number in each cell type of sham and mTBI group. Peach bar represents sham group and turquoise bar represents mTBI group.

Given that the vasculature cells in the initial clustering contain EC (**Sup. Fig. 1C**) and mural cells (**Sup. Fig. 1D-E**), we took the vasculature cells for another round of clustering, aiming for more subtle changes within the vasculature after mTBI. We partitioned the vasculature cells into four clusters (**Fig. 1E**), which can be annotated with known vasculature markers as EC (*Cdh5, Flt1*, and *Pecam1*), PC (*Pdgfrb, Kcnj8*, and *Rgs5*), VSMC (*Acta2* and *Tagln*), and vessel-associated astrocytes (*Aqp4* and *Cldn10*) (**Fig. 1F** and **Sup. Table2**). The clustering outcome was further validated with heatmap analysis of the top 10 markers in each cell type (**Sup. Fig. 2A**). The top 3 marker genes of EC and PC could also be confidently assigned to EC **(Sup. Fig. 2B)** and PC **(Sup. Fig. 2C**), respectively. In addition, the compositions of all four cell types are comparable between sham and mTBI mice (**Fig. 1G**). Considering the abundance of EC and PC over two other cell types and their relevance to microvascular changes, we focus on these EC and PC in the following analysis.

### mTBI induced transcriptional changes in EC and PC

To examine specific genes in EC and PC that may be involved in mTBI induced microvascular injury, we identified differential expression genes (DEGs) between sham and mTBI groups within each vasculature cell cluster (**Sup. Fig. 3**). In the EC-specific DEGs, multiple members of several gene families were enriched, such as heat shock protein family (*Hspa1b, Hspa5, Hspa8, Hspe1*), and ribosomal protein family including the 60S large subunit (*Rpl23a, Rpl36, Rpl39*) as well as the 40S small subunit (*Rps2, Rps24, Rpsa*) (**Fig. 2A** and **Sup. Fig. 4A**). In particular, all four heat shock proteins have >2-fold increases in endothelial expression after mTBI (**Fig. 2A**), which is consistent with previous bulk tissue analysis(Meng et al., 2017). In contrast, members of solute carrier superfamily (*Slc39a10, Slc3a2, Slc40a1, Slc7a1, Slc9a3r2, Slco1a4, Slco1c1*) and transmembrane protein family (*Tmem131, Tmem204, Tmem50a*) were downregulated consistently, except *Tmem252* (**Fig. 2A** and **Sup. Fig. 4A**), indicating a disruption of BBB transport in mTBI. Besides, we also found several EC-specific DEGs that were dramatically downregulated such as *Ddc* and *Ndnf* (***Fig. 2B***), as well as upregulated ones such as *Tmem252* and *Adamts1* (**Fig. 2C**).

**Figure 2.**
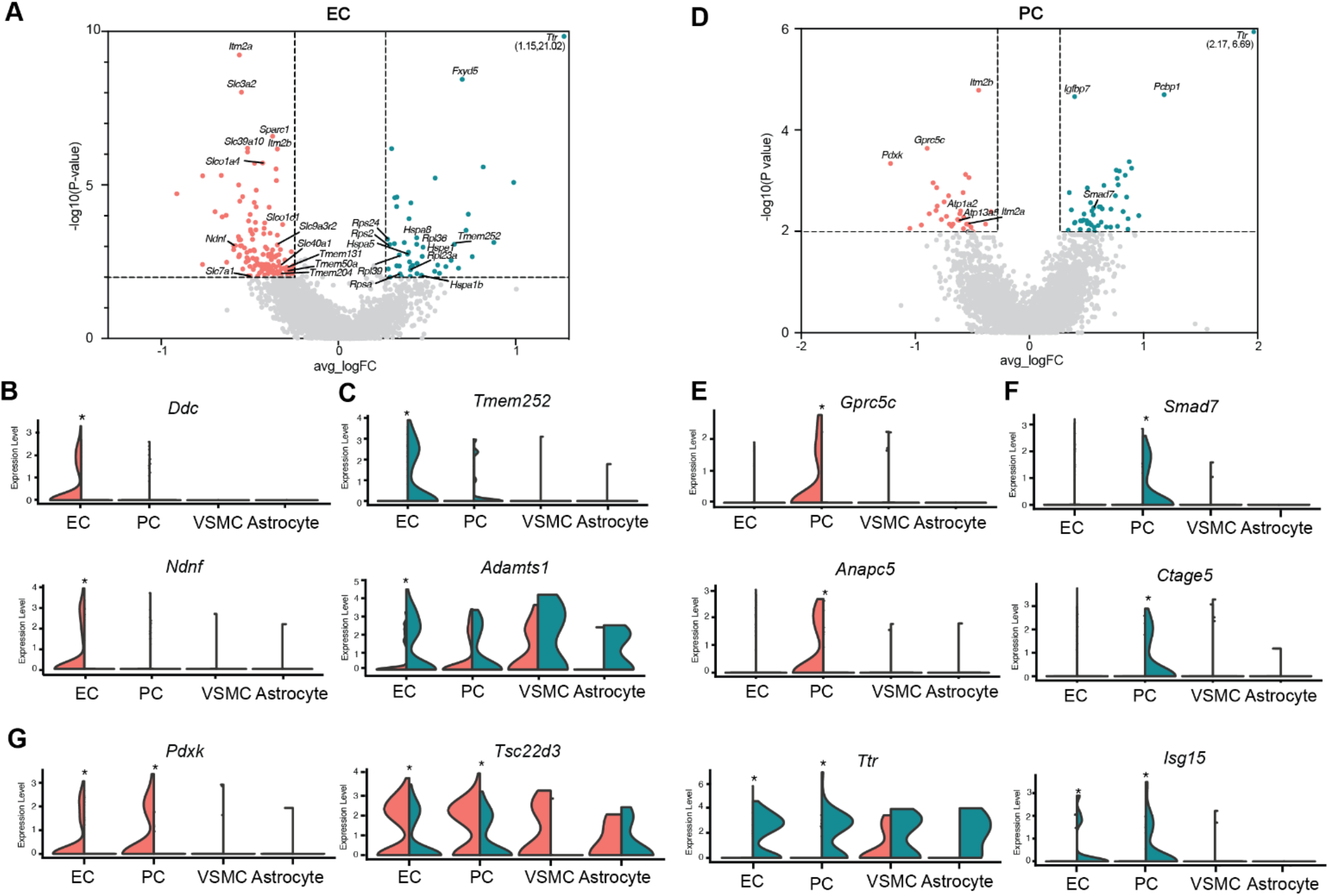
Differentially expressed genes (DEGs) induced by mTBI in EC and PC. **A**, Analysis of differential gene expression was based on sc-RNA seq in EC and was presented as a volcano plot in the mTBI group compared with sham group. **B-C** Split violin plots of the distribution of representative genes in specific cell types. **B** indicated genes specifically decreased in EC and **C**indicated genes specifically increased in EC. **D**, Analysis of differential gene expression was based on sc-RNA seq in PC and was presented as a volcano plot in the mTBI group compared with sham group. The genes in peach are significantly downregulated (−log_10_(p-value) >2 and avg_logFC < −0.25) and the genes in turquoise are significantly upregulated (−log_10_(p-value) >2 and avg_logFC > 0.25). Genes in grey are not significantly changed. **E-G** Split violin plots of the distribution of representative genes in specific cell types. **E**indicated genes specifically decreased in PC, **F** indicated genes specifically increased in PC and **G** indicated genes both changed in EC and PC. Cells from sham group (Peach) are plotted on left, and cells from mTBI group (turquoise) are plotted on right. *P < 0.01.

For the PC-specific DEGs, we found *Atp1a2* and *Atp13a5*, two members of the P-type ATPase superfamily proteins, both decreased by 80% under the mTBI condition (**Fig. 2D** and **Sup. Fig. 4B**). Given that brain PC are directly involved in maintaining BBB function and brain homeostasis(Sweeney et al., 2019), mTBI induced changes in these two ATPase pumps may lead to altered iron and lipid metabolism in the brain. In addition, we also found several PC-specific mTBI responsive genes. For instance, *Gprc5c*, a marker for mural cells(Sheikh et al., 2019), was suppressed by mTBI in PC and could serve as a potential marker for brain injury induced PC alteration (**Fig. 2E**); *Smad7*, a negative regulator of TGF-β signaling(Lee et al., 2017), was significantly increased in mTBI, suggesting that TGF-β signaling may be involved in mTBI-induced BBB dysfunction (**Fig. 2F**).

We also found some DEGs were shared by EC and PC with very similar trends in both cell types (**Sup. Fig. 3B** and **4C**). Notably, transthyretin, encoded by *Ttr*, showed the highest increase in both the EC and PC after mTBI (**Fig. 2G**) as originally reported(Arneson et al., 2018). Interestingly, *Isg15* is an mTBI induced gene found in both EC and PC in this new analysis (**Fig. 2G**). It is linked to intracranial calcifications and viral-induced cognitive impairment in mice(Blank et al., 2016; Hermann and Bogunovic, 2017). In addition, *Pdxk*, which is required for converting Vitamin B6 to its active form pyridoxal-5-phosphate(Elstner et al., 2009), is dramatically reduced in both EC and PC after mTBI (**Fig. 2G**), which may explain the benefits of pyridoxine administration in preclinical TBI models(Kuypers and Hoane, 2010).

### EC and PC specific pathways alteration following mTBI

We performed pathway analysis with KEGG and Gene Ontology databases on the EC and PC specific-DEGs (**Sup. Fig. 3B**). We found EC-specific upregulated DEGs in mTBI were enriched for genes involved in ribosome and protein refolding (**Sup. Fig. 5A**, red), which is consistent with the highly upregulated heat shock proteins in brain injury(Kim et al., 2020). Our analysis also highlighted that downregulated EC-specific DEGs were significantly enriched in pathways important for extracellular matrix (ECM) organization and modulating neural projection (**Sup. Fig. 5A**, blue). In addition, downregulated DEGs were enriched in transport-related pathways such as anion transport and lipid transport **(Sup. Fig. 5A**, blue), suggesting the disruption of transport function in BBB. PC-specific pathway analysis demonstrated that upregulated DEGs are involved in leukocyte trans-endothelial migration, GTPase activity, and oxidoreductase activity (**Sup. Fig. 5B**, red). In contrast, PC-specific downregulated DEGs are involved in the dendrite, mitochondrial membrane, and ATP metabolic process (**Sup. Fig. 5B**, blue).

### Aging-associated transcriptional changes in EC and PC

Since microvascular pathology is also common in aging(Montagne et al., 2015), we applied our analysis workflow on the aging scRNA-seq dataset(Ximerakis et al., 2019). UMAP was able to separate cell into clusters, which were mapped to 11 cell types (astrocyte, oligodendrocyte, OligoPC, microglia, macrophage, vascular cell, ependymal cell, and 4 neuron subtypes) (**Fig. 3A**). Vasculature cell cluster contains EC (**Sup. Fig. 6A**) and mural cells (**Sup. Fig. 6B** and **6C**). Therefore, we took all vasculature cells for a second round of clustering, which revealed 6 subclusters including EC, PC1, PC2, VSMC, vessel-associated astrocyte, and Hemoglobin-expressing EC (**Fig. 3B**). Annotation with known vasculature markers helped to resolve EC (*Cdh5, Cldn5, Flt1*, and *Pecam1*), PC (*Pdgfrb, Kcnj8*, and *Rgs5*), VSMC (*Acta2* and *Tagln*), and vessel-associated astrocyte (*Aqp4* and *Cldn10*) (**Fig. 3C** and **Sup. Table 4**). Unbiased clustering for vascular cells indicated two types of pericytes (PC1 and PC2), which expressed general PC markers (*Kcnj8* and *Art3*). We also found some PC markers specifically expressed in subtypes of PC (Vanlandewijck et al., 2018), such as *Trpc3* for PC1, and *Ggt1* for PC2 (**Sup. Fig. 6D**).

**Figure 3.**
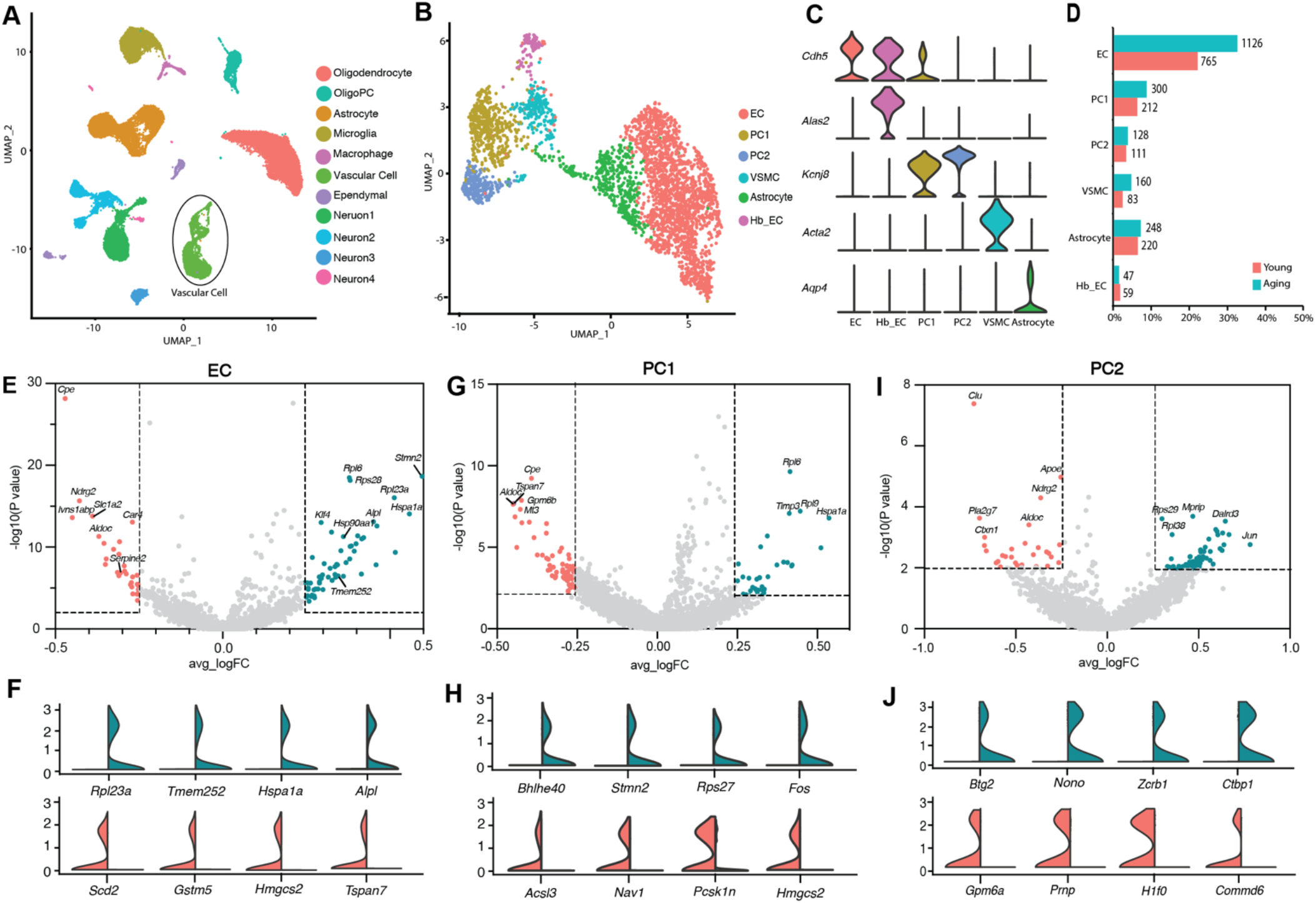
Secondary analysis of cerebrovascular changes in aging scRNA-seq dataset. **A**, UMAP showing cell clusters which were color coded and annotated based on their transcriptional profile identities. Vascular cells are marked by black line. **B**, UMAP visualization of 3459 vascular cell transcriptomes (1450 from young mouse brain and 2009 from aging mouse brain). UMAP showing cell clusters which were color coded and annotated based on their transcriptional profile identities. **C**, Violin plots showing gene markers that distinguish across vasculature cells. Genes are colored by cell types. **D**, Bar plot showing the percentage of cells associated with EC, Hb_EC, PC1, PC2, VSMC, and Astrocyte in both young and aging group. The number besides the bar shows cell number in each cell type of young and aging group. Peach bar represents young group and turquoise bar represents aging group. Hb_EC: Hemoglobin_EC. **E**, Analysis of DEGs was based on aging sc-RNA seq in EC and was presented as a volcano plot in the aging mice compared with young mice. **F**, Split violin plots showing the distribution of representative genes in EC. **G**, Analysis of DEGs was based on aging sc-RNA seq in PC1 and was presented as a volcano plot in the aging mice compared with young mice. **H**, Split violin plots showing the distribution of representative genes in PC1. **I**, Analysis of DEGs was based on aging sc-RNA seq in PC2 and was presented as a volcano plot in the aging mice compared with young mice. **J**, Split violin plots showing the distribution of representative genes in PC2.

Moreover, it reveals approximately 60% of vascular cells are EC, 20% are PC, and 7% are VSMC, which shared a similar proportion of vascular cell population with mTBI dataset (**Fig. 3D**).

To identify the molecular signatures of aging-induced microvascular injury, we focused on DEGs of EC, PC1, and PC2 in the aging dataset. There are multiple DEGs in EC (**Fig. 3E** and **Sup. Table5**); yet, the changes of multiple gene families in aged EC is in line with those from the mTBI dataset. For example, heat shock protein family (*Hspa1a, Hsp90aa1*) and ribosomal protein family (*Rpl6, Rps28, Rpl23a, Rpl36-ps3*) were upregulated, while the solute carrier superfamily (*Slco2b1, Slc38a5, Slc16a4, Slc1a2*) were downregulated in EC (**Fig. 3E** and **F**). In addition, we found two PC subtypes varied considerably in aging-responsive genes with only a small overlap (**Sup. Fig. 6E**), suggesting that they are functionally different in aging. Among the DEGs in PC1 (**Fig. 3G**), *Stmn2*, a stathmin phosphoprotein being implicated in AD(Meyer et al., 2019), Parkinson’s disease(Wang et al., 2019) and amyotrophic lateral sclerosis(Klim et al., 2019), was dramatically increased in aging (**Fig. 3H**). On the other hand, in PC2 DEGs (**Fig. 3I**), *Zcrb1*, a gene previously known to be upregulated in aging microglia(Orre et al., 2014), was upregulated; while *Gpm6a*, a transmembrane protein plays a key role in cognitive function(Gregor et al., 2014), was significantly decreased in PC2 (**Fig. 3J**).

### Identification of the common molecular signatures of microvascular injury

To explore the common molecular signature of microvascular injury in EC and PC, we compared DEGs from the mTBI dataset with DEGs from the aging dataset, respectively. The venn diagrams indicated six shared microvascular injury induced molecular signatures in EC, including three increased signatures (*Tmem252, Adamts1*, and *Rpl23a)*, and three decreased signatures *(Car4, Ndnf*, and *Serpine2)* **(Fig. 4A)**, and two shared molecular signatures in PC2, including increased *Rps29* and decreased *Sepp1 (***Fig. 4B**). However, there is no common molecular signature between PC1 in mTBI and aging (data not shown). To facilitate the understanding of shared pathways, meta-analysis of DEGs from the mTBI dataset as well as the aging dataset was used through Metascape(Zhou et al., 2019). **Fig. 4C** and **4D** depict the enriched biological pathways from KEGG and Gene Ontology across two datasets. We observed that aging and mTBI induced microvascular injury in EC is closely related to vascular development and changes of the ECM (**Fig. 4C**), and the common molecular signatures including *Adamts1, Serpine2*, and *Ndnf* are involved in the above shared pathways (**Fig. 4C**). In addition, decreased endothelial *Car4* expression was associated with the monocarboxylic acid transport pathway(**Fig. 4C**), which is involved in many neuropathological disorders, including PD and AD(Prins and Giza, 2006)(Halestrap, 2013), suggesting monocarboxylic acid transport dysfunction in microvascular injury. Meanwhile, one of the common molecular signatures in PC, *Rps29*, is involved in protein biosynthesis resulting from the translation of messenger RNA (**Fig. 4D**).

**Figure 4.**
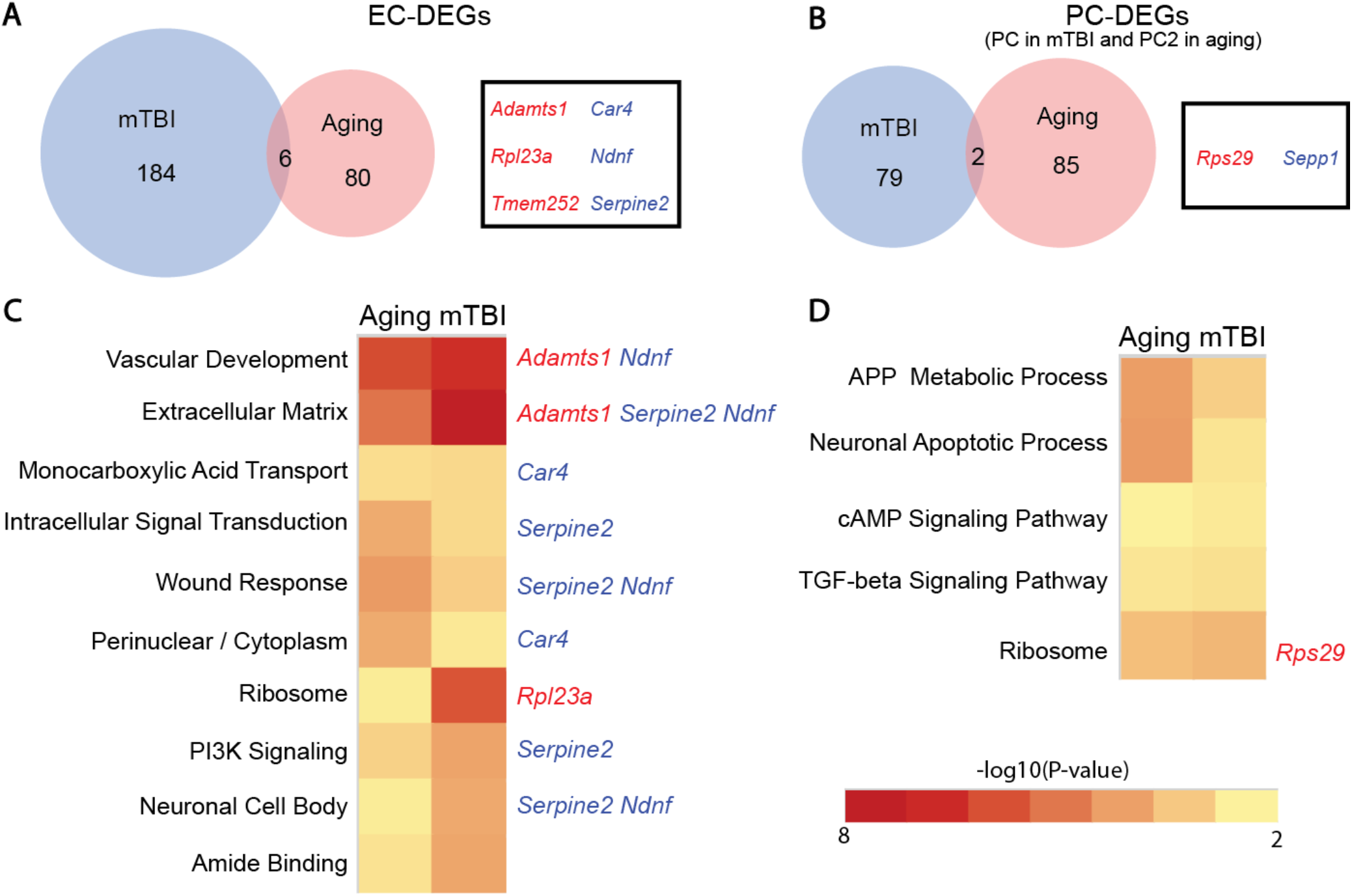
Identification common molecular signatures of microvascular injury. **A-B**, Venn Plot indicated that overlaid DEGs in EC (**A**) and PC (**B**) between mTBI dataset and aging dataset. **C-D**, the heatmap indicated the −log_10_(P-value) of aging and mTBI dataset in shared pathways in EC (**C)** and PC **(D)**. The left side of heatmap shows the corresponding pathways, and the right side of heatmap indicate the shared DEGs in corresponding pathways. The increased genes are marked by red, and the decreased genes are marked by blue.

Taken together, we have developed a workflow to understand the transcriptional programs of cerebrovascular cells during microvascular injury and aging. The secondary analysis on vascular cells uniquely points to cell-type specific changes in gene expression and biological pathways that might bring novel insights into the function of EC and PC in microvessel injuries in mTBI, aging, and beyond.

## Discussion

Given that gene expression programs and gene synergy determine the cellular function to respond to external signals and to carry out complex cellular activities, high-throughput sequencing technologies are now in place to discover gene expression programs and explore the biological mechanism that they govern. Existing genomic profiling studies for cerebrovascular cells use the marker-based method to isolate a single type of vasculature cells for further sequencing(Munji et al., 2019). However, distinct cell types in BBB have their own unique functions and perform highly coordinated actions to keep the central nervous system (CNS) homeostasis(Zhao et al., 2015); therefore, it is important to accurately distinguish different vascular components in the brain to explore the potential function in BBB disruption. ScRNA-seq is a good method to capture cellular transcriptomes profiling in an unbiased manner, allowing investigation of individual cells’ expression signature. In this study, we first investigated cerebrovascular injury through secondary scRNA-seq analysis for vascular cells and illustrated transcriptional profiling in cerebrovascular changes in the mTBI and aging mice to define the underlying molecular signatures of vascular impairment which is common among distinct pathological conditions. The transcriptome patterns of common molecular signatures depict gene programs that may be involved in BBB dysfunction and provide new insights for the possible vascular mechanism to post-mTBI neurodegenerative disorders such as AD.

It is striking that three common molecular signatures are the component of ECM in the brain, including increased extracellular protease *Adamts1*, decreased extracellular protease inhibitor *Serpine2*, and decreased neuron-derived neurotrophic factor (*Ndnf*) under mTBI condition. *Adamts1*, a risk gene for late-onset Alzheimer’s Disease(Kunkle et al., 2019), is required for ECM remodeling(Brown et al., 2006) and the regulation of artery shear stress(Kalucka et al., 2020). Additionally, many of the extracellular proteases are increased along vessels in BBB dysfunction(Munji et al., 2019). *Ndnf* is also found the specific expression in EC, but barely expressed after mTBI. *Ndnf* is secreted from EC and accelerates EC function and ischemia-induced angiogenesis, leading to blood flow recovery and tissue perfusion(Ohashi et al., 2014). Besides, *Ndnf* is important to protect hippocampal neurons from the toxicity of beta-amyloid peptide(Kuang et al., 2010). Given that the composition of the ECM is altered after BBB disruption and directly affects the progression of neurodegenerative disease(Lendahl et al., 2019), the disruption of ECM may be an important link between mTBI induced BBB dysfunction and secondary neurodegeneration.

Transport dysfunction is one of the hallmarks of BBB disruption(Zhao et al., 2015). In our analysis, decreased expression of BBB transporters was observed in cerebrovascular injury associated with both mTBI and aging. The disrupt of transporter after TBI strongly suggests BBB dysfunction, particularly in clearing metabolites, which may also correlate to neurodegenerative changes similarly observed in AD(Sweeney et al., 2019). Another common molecular signature in a subtype of PC (PC2), *Sepp1*, is also decreased in microvascular injury. *Sepp1* is important in the transport of selenium via apolipoprotein E recaptor-2 (apoER2). Mice with Sepp1 deficiency develop brain injury with neurological dysfunction, which can be attributed to the impairment of Sepp1-apoER2 interaction at the BBB(Burk et al., 2014). Interestingly, a transmembrane protein, *Tmem252*, with unknown function is highly and specifically expressed in EC under mTBI condition and aging process but almost absent in normal condition. Coincidentally, *Tmem252* is also dramatically upregulated in stroke (Zeng et al., 2019) and LPS-induced vascular inflammation(Chen et al., 2020), suggesting that *Tmem252* may be a common regulator of cerebrovascular injury.

PC dysfunction has been reported in many pathologic conditions, such as mTBI, stroke, and AD(Sweeney et al., 2018). PC are required for the integration of BBB, and PC loss can cause impairment of PC-EC crosstalk, leading to BBB dysfunction(Bhowmick et al., 2019). The change of shear stress after mTBI could inhibit TGF-*β* activation, which is required PC-EC interaction(Sweeney et al., 2016). Thus, the sharp rise of *Smad7* in PC after mTBI implicated the weak interaction between EC and PC and the change of shear stress, leading to BBB dysfunction. This suggests that PC changes may contribute to BBB dysfunction and the increased risk of AD after mTBI. The fact that only one shared DEG is found between PC in mTBI and aging suggests that PC is probably heterogenous and undergoes differential changes between different subtypes. The two potential subtypes discussed here are perhaps just the tip of the iceberg, more in-depth studies are required to unveil the mystery.

Additionally, a small fraction of choroid plexus vulnerable to mTBI was detected in the hippocampus, a ventricle-adjacent region. Choroid plexus cells were abundant in mTBI conditions (4% from mTBI group and 0% from Sham group). The choroid plexus, as a responder to injury, is the site of the blood-cerebrospinal fluid barrier(Xiang et al., 2017). It has been reported that external stimuli such as injury can cause choroid plexus epithelial cell proliferation (Xiang et al., 2017). Moreover, the alteration of the choroid plexus has been observed in AD(Kaur et al., 2016). Thus, this large shift in the number of choroid plexus provides new insight into the mechanism between mTBI and AD. However, many external factors such as tissue preparation, sensitivity to tissue dissociation, sequencing procedure may lead to the differences between actual and estimated percentages for each cell type, and in-depth studies are needed in the future.

## Supplementary Data

**Sup. Fig. 1.**
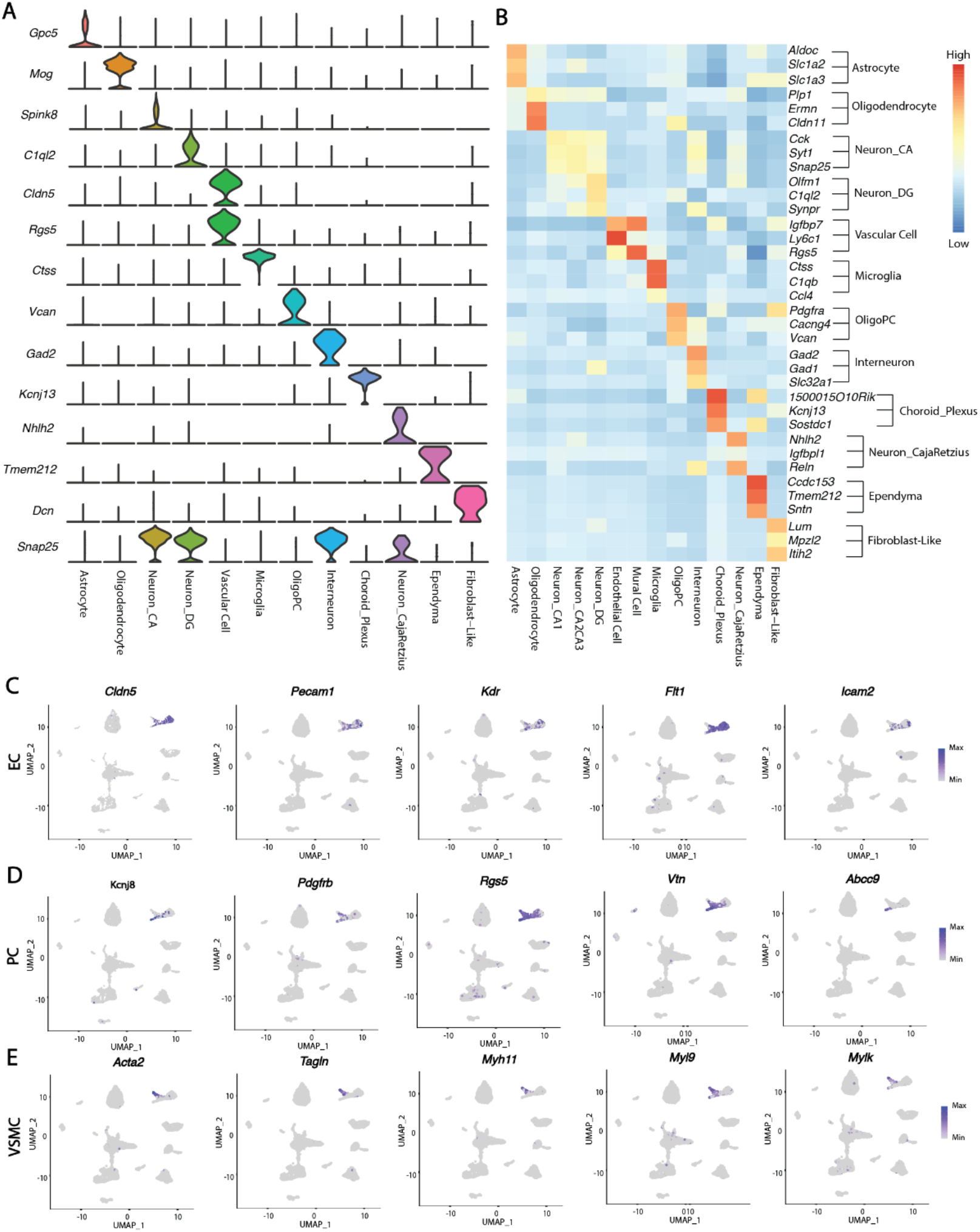
Identification of cell types in mTBI scRNA-seq dataset. **A**, Violin plots showing gene markers that distinguish across and within brain cell types. Genes are colored by cell types. **B**, Gene-expression heatmap of the top 3 marker genes in DropViz’s dataset for each cluster. The cell type in rows shows the cell clusters in DropViz’s dataset. Color scale: red, high expression; blue: low expression. **C**, UMAP visualization of vascular cell population showing the expression of representative EC markers *Cldn5, Pecam1, Kdr, Flt1*, and *Icam2* respectively. **D**, UMAP visualization of vascular cell population showing the expression of representative PC markers *Kcnj8, Pdgfrb, Rgs5, Vtn*, and *Abcc9* respectively. **E** UMAP visualization of vascular cell population showing the expression of representative VSMC markers *Acta2, Tagln, Myh11, Myl9*, and *Mylk* respectively.

**Sup. Fig. 2.**
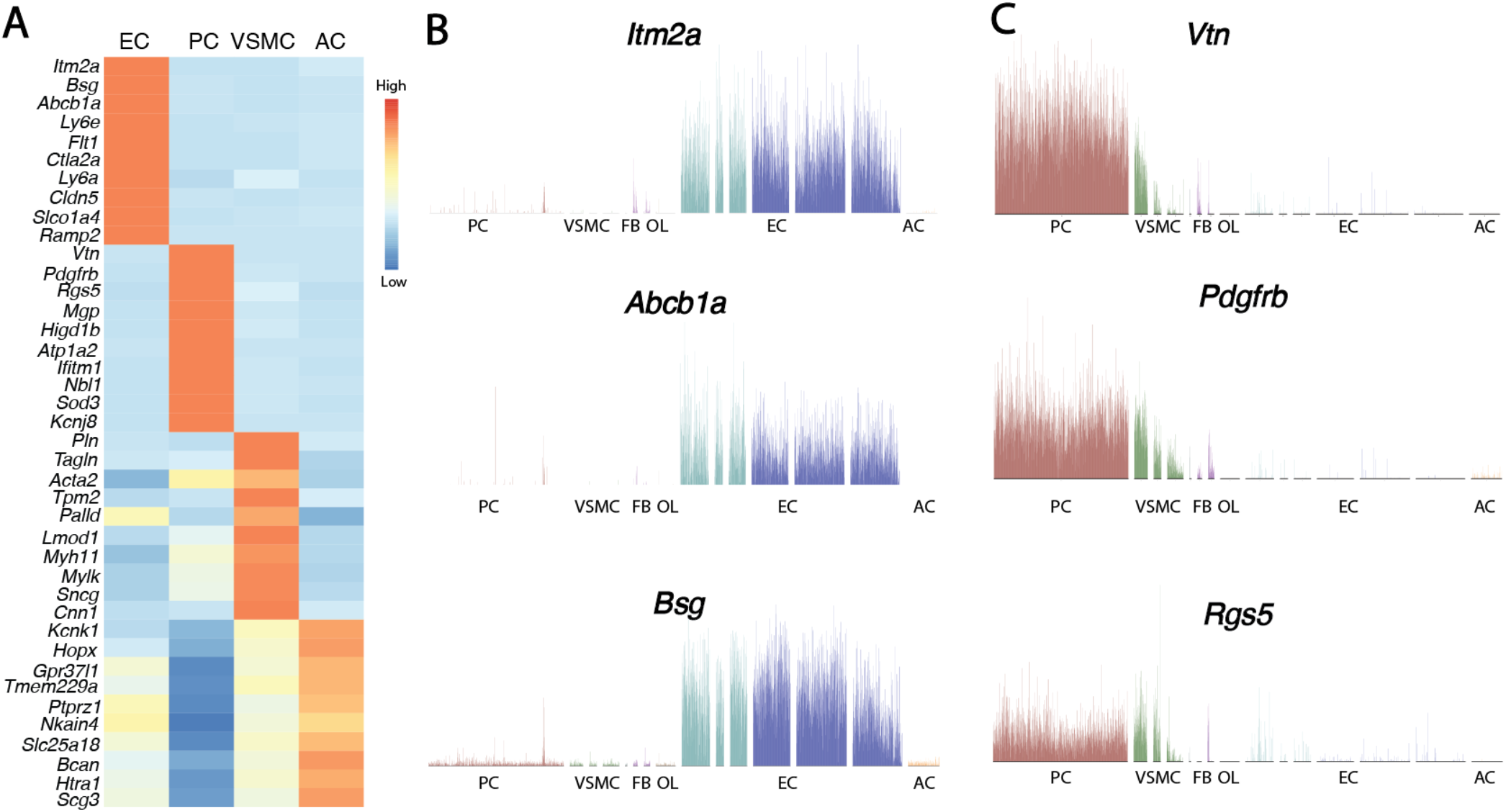
Identification of cell types in vasculature cells in mTBI dataset. **A**, Gene-expression heatmap of the top 10 marker genes in vasculature cells for each cluster. Color scale: red, high expression; blue: low expression. AC: Astrocyte. **B**, Bar plots showing top 3 EC marker expression pattern in brain vasculature dataset. **C**, Bar plots showing top 3 PC marker expression pattern in brain vasculature dataset.

**Sup. Fig. 3.**
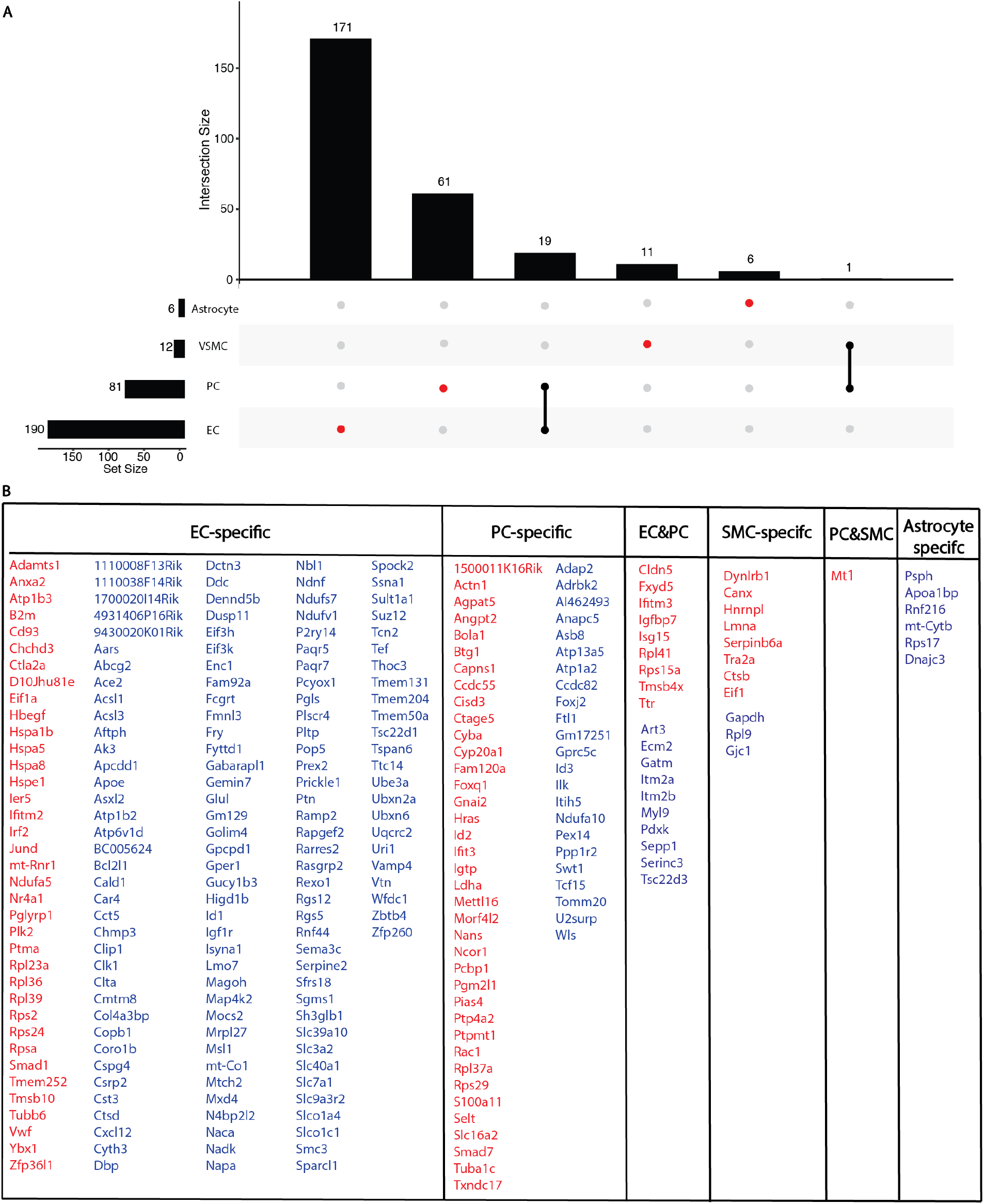
Identification of cell type specific DEGs. **A**, DEGs unique to a cell type are indicated in red and those shared between >= 2 cell types are indicated by black dots. The histogram above each plot indicated the DEG counts for each category. The bar plot besides the histogram indicated the DEG counts for each cell type. **B**, A table showing the genes in each category.

**Sup. Fig. 4.**
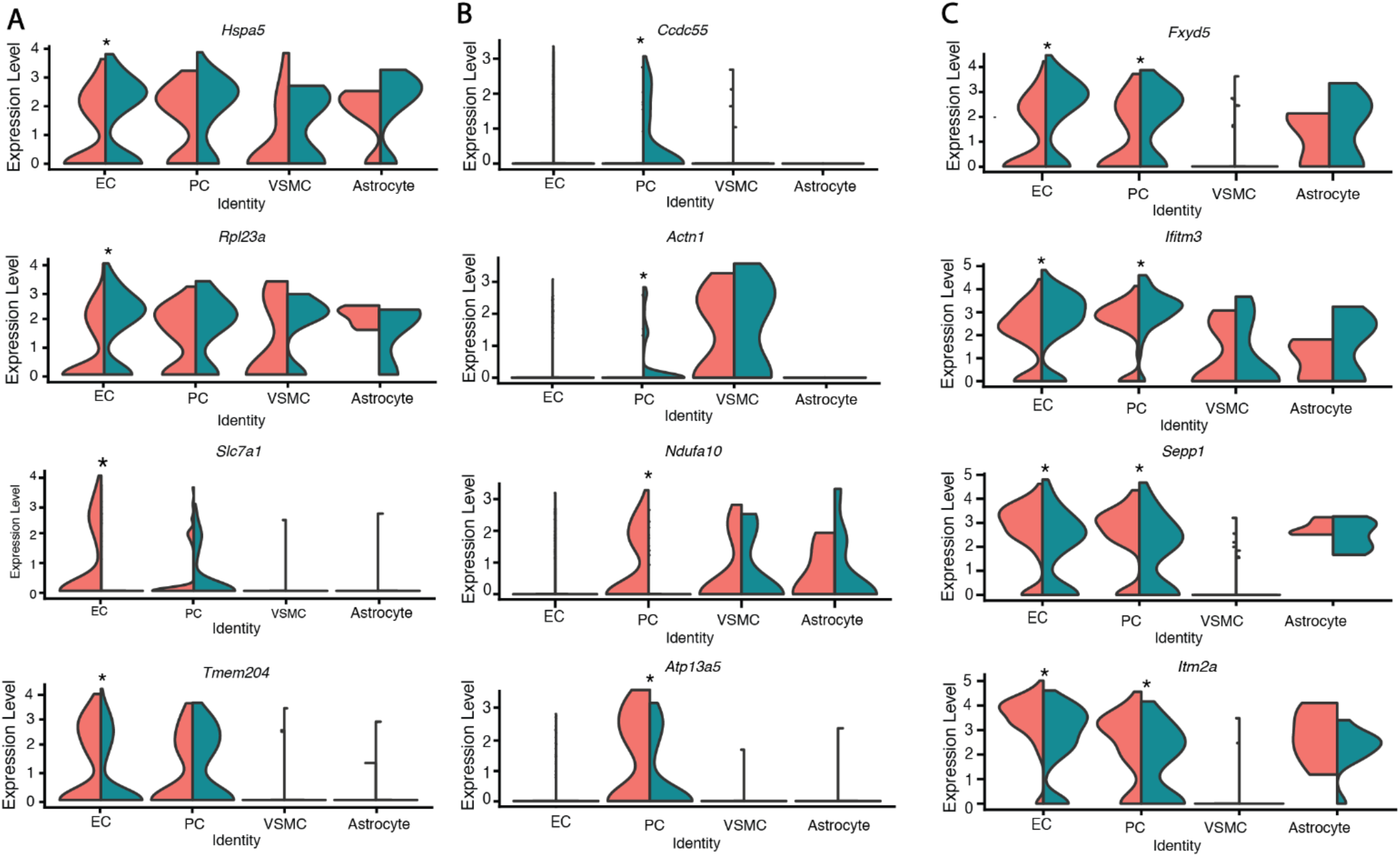
Representative DEGs induced by mTBI in EC and PC. Split violin plots of the distribution of representative genes in specific cell types. **A** indicated genes specifically changed in EC, **B** indicated genes specifically changed in PC, and **C** indicated genes both changed in EC and PC. Cells from sham group (Peach) are plotted on left, and cells from mTBI group (turquoise) are plotted on right. *P < 0.01.

**Sup. Fig. 5.**
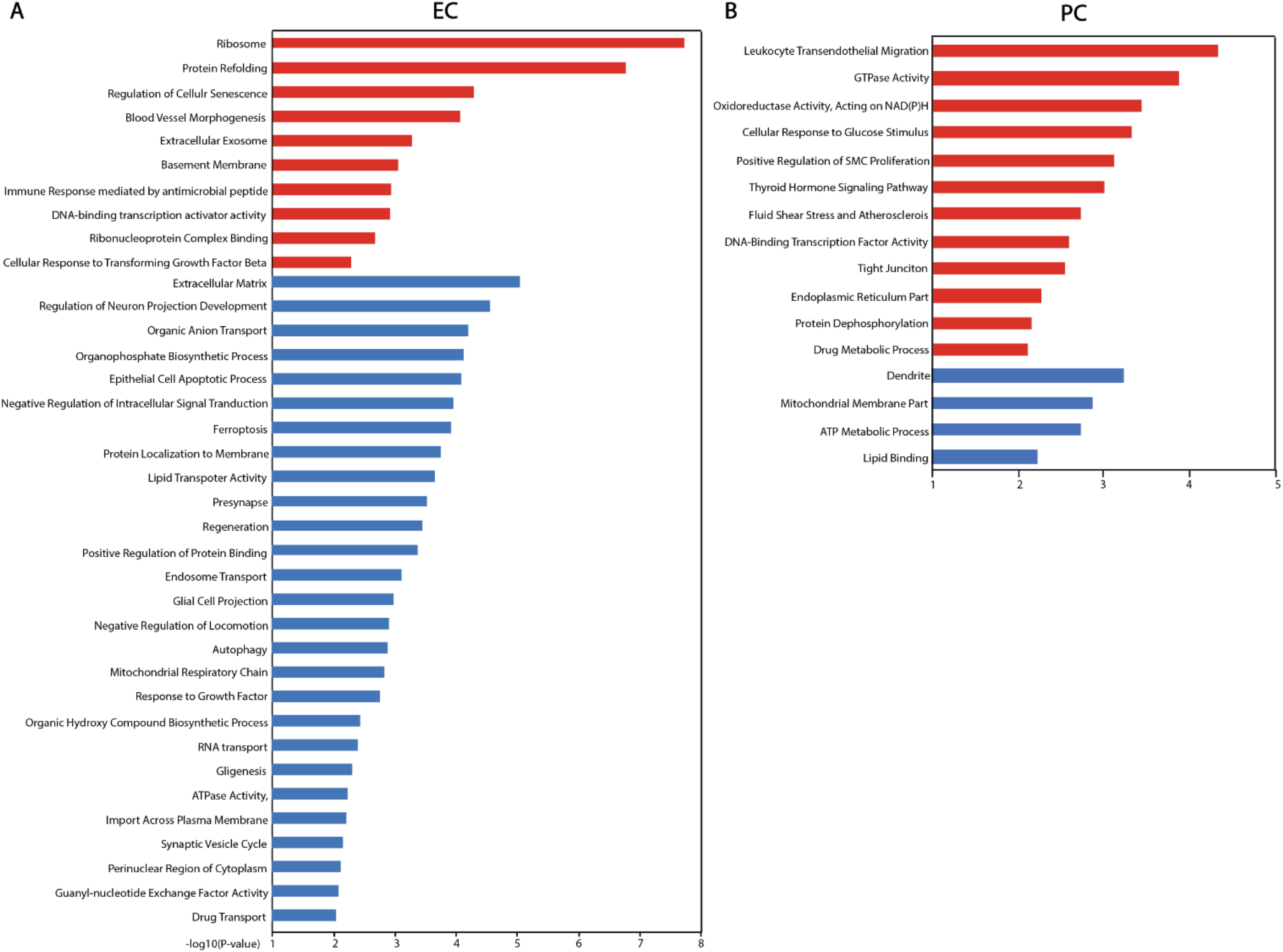
Pathways analysis on EC and PC specific DEGs. Pathway analysis was used to identify the most significant upregulated and downregulation GO terms and KEGG pathways based on the EC or PC specific DEGs under mTBI condition. **A**, Pathways analysis on EC. Red bar indicated the pathways enriched by upregulated EC specific DEGs, and blue bar indicated the pathways enriched by downregulated EC specific DEGs. **B**, Pathways analysis on PC. Red bar indicated the pathways enriched by upregulated PC specific DEGs, and blue bar indicated the pathways enriched by downregulated PC specific DEGs. The pathways with −log_10_(P-value) > 2 are presented.

**Sup. Fig. 6.**
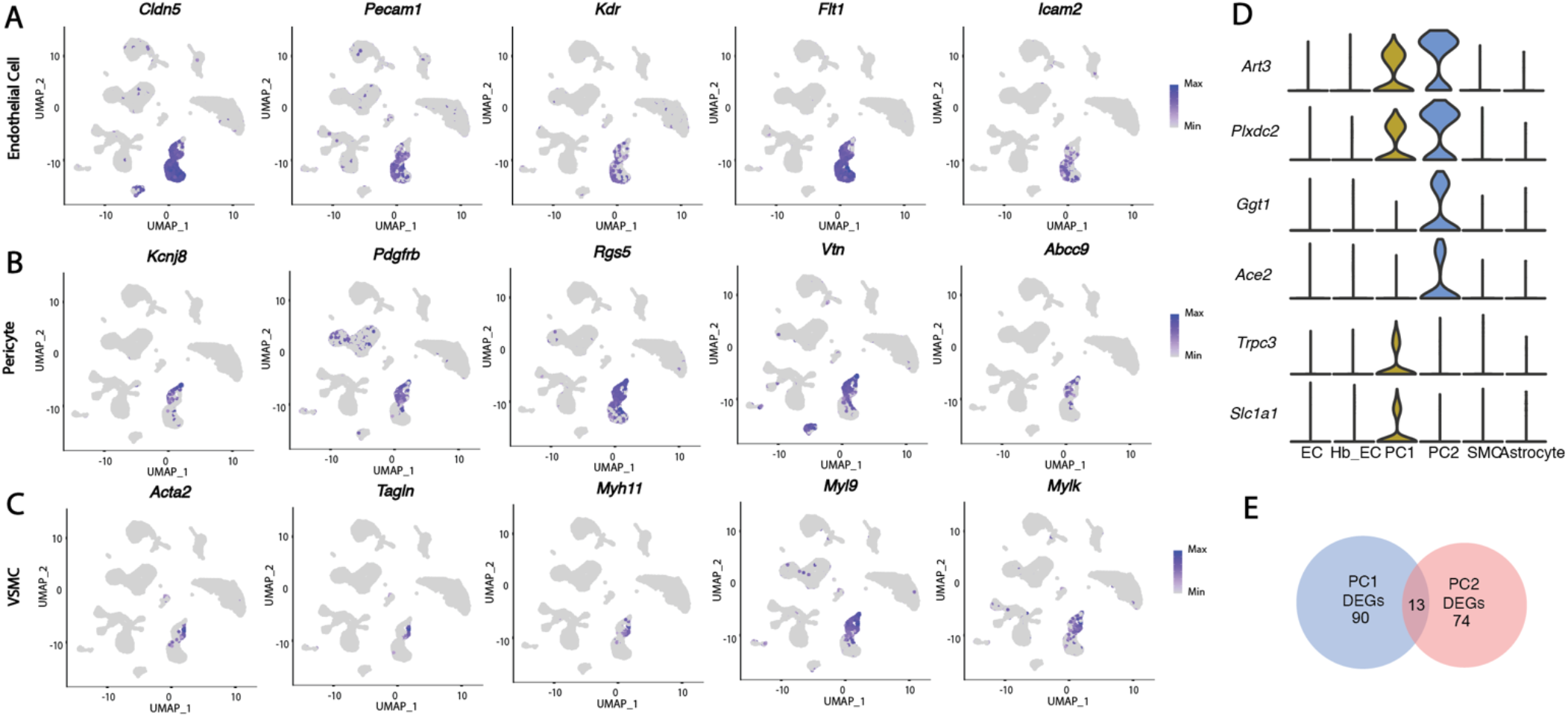
Identification of cell types in vasculature cells in aging dataset. **A**, UMAP visualization of vascular cell population showing the expression of representative EC markers *Cldn5, Pecam1, Kdr, Flt1*, and *Icam2* respectively. **B**, UMAP visualization of vascular cell population showing the expression of representative PC markers *Kcnj8, Pdgfrb, Rgs5, Vtn*, and *Abcc9* respectively. **C**, UMAP visualization of vascular cell population showing the expression of representative VSMC markers *Acta2, Tagln, Myh11, Myl9*, and *Mylk* respectively. **D**, Violin plots showing gene markers that distinguish across and within vascular cell types. Genes are colored by cell types. **E**, Venn Plot indicated that overlaid DEGs between PC1 and PC2 from aging dataset.

